# A Comparative Evaluation of Structural MRI Foundation Models for Age, Sex, and Body-Mass Index Predictions

**DOI:** 10.64898/2026.05.15.725427

**Authors:** Arel Encin, Asa Gilmore, Ariel Rokem, Erin Dickie, Tristan Glatard

## Abstract

Foundation models pre-trained on large neuroimaging datasets offer a promising approach to overcome the limited sample sizes typical of clinical imaging studies, yet their generalization across diverse populations remains unclear. We present the first systematic benchmark of four publicly available structural MRI foundation models: AnatCL, BrainIAC, 3D-Neuro-SimCLR, and SwinBrain. Using T1-weighted MRIs from the Parkinson’s Progression Markers Initiative (PPMI), Healthy Brain Network (HBN), and Nathan Kline Institute (NKI) datasets, we evaluate these models on sex classification, brain age prediction, and body mass index prediction, comparing against models trained from FreeSurfer-derived cortical thickness and cortical surface area features. Submitted models are evaluated using a standardized frozen feature probing framework. The evaluation methods are available in BrainFM-Bench, a living benchmark for structural brain MRI foundation models hosted on GitHub, where new models can be added through pull requests. Although some foundation models outperformed FreeSurfer on particular tasks and datasets, 3D-Neuro-SimCLR and AnatCL outperformed the baselines overall, with 3D-Neuro-SimCLR demonstrating the most consistent performance (with the notable exception of HBN sex classification). The remaining models did not consistently outperform the baselines, indicating that the advantage of learned representations over morphometric features was not consistent across datasets and tasks for these models. In addition, cross-model feature correlation analysis reveals that foundation model representations correlate differently with traditional cortical measurements. These findings position structural MRI foundation models, particularly 3D-Neuro-SimCLR and AnatCL, as promising avenues to boost the performance of predictive models in neuroimaging.

## 1 Introduction

Foundation models have the potential to address the sample size limitations of neuroimaging studies through pre-training on large databases. Nonetheless, for these models to generalize well, their pre-training must adequately reflect the broad range of populations encountered in practice, which remains a significant challenge. This paper aims to systematically evaluate publicly available structural MRI brain foundation models on phenotypic prediction tasks. Using T1-weighted (T1w) images from the Parkinson’s Progression Markers Initiative (PPMI) [12], an older-adult cohort, the Nathan Kline Institute (NKI) [14], a community sample spanning a broad adult age range, and the Healthy Brain Network (HBN) [1], a children and adolescent cohort, we evaluate four structural brain MRI foundation models: AnatCL, BrainIAC, 3D-Neuro-SimCLR, and SwinBrain. We compared their performance on sex classification, brain age prediction, and body mass index (BMI) regression with that of models based on FreeSurfer morphometric features. Additionally, we explored whether the representations learned by these foundation models provide complementary information to standard FreeSurfer-derived measures. Our main contributions are as follows: (1) we provide the first benchmarking analysis of structural brain MRI foundation models across sex classification, brain age prediction, and BMI regression, and (2) we examine the relationship between foundation model representations and traditional FreeSurfer-derived features in terms of feature relevance; and (3) we introduce BrainFMBench, a living benchmark that enables standardized, automated evaluation of structural brain MRI foundation models through frozen feature probing on downstream tasks.

## 2 Materials & Methods

### 2.1 Overall Benchmarking Approach

All foundation models were used as frozen feature extractors: each model produced a fixed-length representation per scan, with no fine-tuning of encoder weights. We uniformly applied a Random Forest (RF) classifier for sex classification and a Random Forest regressor for brain age and BMI prediction to all feature sets. RF hyperparameters were set to conservative values without task-specific tuning to ensure a consistent evaluation across all models. We used an RF probe rather than a tuned or task-specific model because our aim was to compare representations rather than to maximize downstream performance: a single probe applied identically to every feature set isolates differences attributable to the representations themselves and handles non-linear structure without per model tuning. For classification, we used *n*_estimators_ = 200, max_depth = 6, min_samples_split = 5, 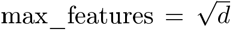 (where *d* is the feature dimensionality), and balanced class weights. For regression, we used the same settings except max_features = *d* (all features) and no class weighting. Features were z-score standardized using statistics computed from the training set only. For each task, we performed a 90/10 train-test split (stratified by the target label for the classification task), followed by 5-fold cross-validation on the training portion (stratified for classification), repeated across five random seeds to assess stability. Classification performance was evaluated using balanced accuracy, while regression performance was measured using the mean absolute error (MAE). 3D-Neuro-SimCLR and BrainIAC were excluded from the PPMI evaluation, as both models included PPMI data in their pretraining datasets.

### 2.2 Structural Brain MRI Foundation Models

*AnatCL* [3] is a weakly contrastive foundation model for structural brain MRI that extends standard contrastive learning to continuous labels by computing a degree of positiveness between samples. It combines age-based alignment with anatomical features (cortical thickness, gray matter volume, surface area) derived from atlas parcellations. The framework offers two descriptors, both evaluated in this paper: a *local* descriptor that captures region-specific variation via cross-region similarity across ROIs, and a *global* descriptor that aggregates features brain-wide via cross-measure similarity. The model is pre-trained on 3,984 healthy subjects from OpenBHB [5] using a ResNet-18 encoder that produces 512-dimensional feature vectors. The authors provided five fold checkpoints for each descriptor; as recommended by the authors, we extracted features from all five folds and then averaged them.

*BrainIAC* [19] is a self-supervised foundation model for multiparametric brain MRI trained via SimCLR contrastive learning on 48,965 scans spanning multiple sequences (T1, T2, T1CE, FLAIR) and ten clinical conditions across 34 datasets. The framework employs NT-Xent loss to maximize agreement between augmented views of the same image while repelling negative pairs. We used the publicly released Vision Transformer (ViT-B/16) encoder, which produces 768-dimensional representations.

*3D-Neuro-SimCLR* [11] is a self-supervised model trained via SimCLR contrastive learning on 44,958 T1-weighted scans from 11 public datasets, using a ResNet-18 encoder producing 512-dimensional representations.

*SwinBrain* [18] is a self-supervised foundation model for clinical multi-contrast brain MRI trained using Swin UNETR with a cross-contrast context recovery task on 75,861 scans across 14 MRI scanners. The framework concatenates the T1-weighted, T2-weighted, and FLAIR sequences into a 3-channel volume and learns to recover masked regions using both reconstruction and contrastive losses, producing 384-dimensional representations. Because our evaluation datasets contain only T1-weighted images, we duplicated the T1w volume across all three input channels.

### 2.3 Baseline model

#### FreeSurfer

To investigate feature relevance, we obtained cortical thickness and surface area values from the Desikan-Killiany (aparc) [4] and Schaefer 400 [15] parcellations using FreeSurfer [6] for all participants across the HBN, NKI, and PPMI datasets. For the HBN and NKI datasets, FreeSurfer features were obtained from the Reproducible Brain Charts (RBC) initiative [16], processed with FreeSurfer v6.0.1. For the PPMI dataset, FreeSurfer features were obtained from FreeSurfer v7.3.2 [12]. Following the approach of Modabbernia et al. [13], we used surface area and cortical thickness as features for the downstream evaluations.

### 2.4 Datasets and Preprocessing

Structural T1-weighted MRI scans were obtained from PPMI (894 subjects: 555 male, mean age 62.48 *±* 9.99 years), HBN (1,000 subjects: 519 male, mean age 11.33 *±* 3.77 years; BMI available for 963 subjects, mean 19.7 *±* 5.1), and NKI (958 subjects: 357 male, mean age 41.99 *±* 20.22 years; BMI available for 956 subjects, mean 26.5 *±* 5.8). Sex classification, age prediction, and BMI regression (HBN and NKI only) were evaluated. AnatCL images were preprocessed with CAT12 [7] (modulated warped gray matter maps, 1.5 mm isotropic). All other models used TurboPrep^8^ (N4 bias correction [20], skull stripping [8], MNI registration [2], intensity normalization [17], 1 mm isotropic), with model-specific input dimensions and z-score normalization. To verify that TurboPrep did not bias results in favor of 3D-Neuro-SimCLR, BrainIAC and SwinBrain were also evaluated under BrainIAC’s training pipeline (N4, resampling, rigid MNI registration, HD-BET [10]) and a variant using the BrainIAC repository’s reference template. 3D-Neuro-SimCLR and BrainIAC were excluded from PPMI as both were pretrained on it. FreeSurfer baselines used the aparc atlas for PPMI (v7.3.2 [12], 140 features) and both aparc and Schaefer 400-parcel atlases for HBN and NKI (v6.0.1 via RBC [16], 136 and 800 features respectively).

## 3 Results

We implemented our core benchmark as a living benchmark hosted on GitHub https://github.com/epics-lab/BrainFMBench, enabling external users to contribute new models through pull requests. Additional analysis scripts used for supplementary experiments are provided separately at https://github.com/epics-lab/additional-evaluation-brainfm.

### 3.1 3D-Neuro-SimCLR and AnatCL outperformed baseline overall

Overall, AnatCL and 3D-Neuro-SimCLR stood out as the most accurate models, outperforming the FreeSurfer baselines across most tasks and datasets (Fig. 1). AnatCL significantly outperformed FreeSurfer baselines on age prediction, but its performance was less consistent for sex classification and BMI regression. 3D-Neuro-SimCLR significantly outperformed both baselines on every HBN and NKI task except HBN sex classification. The remaining models did not consistently outperform the FreeSurfer baselines, indicating that the advantage of learned representations over morphometric features was not consistent across datasets and tasks for these models.

**Fig. 1:**
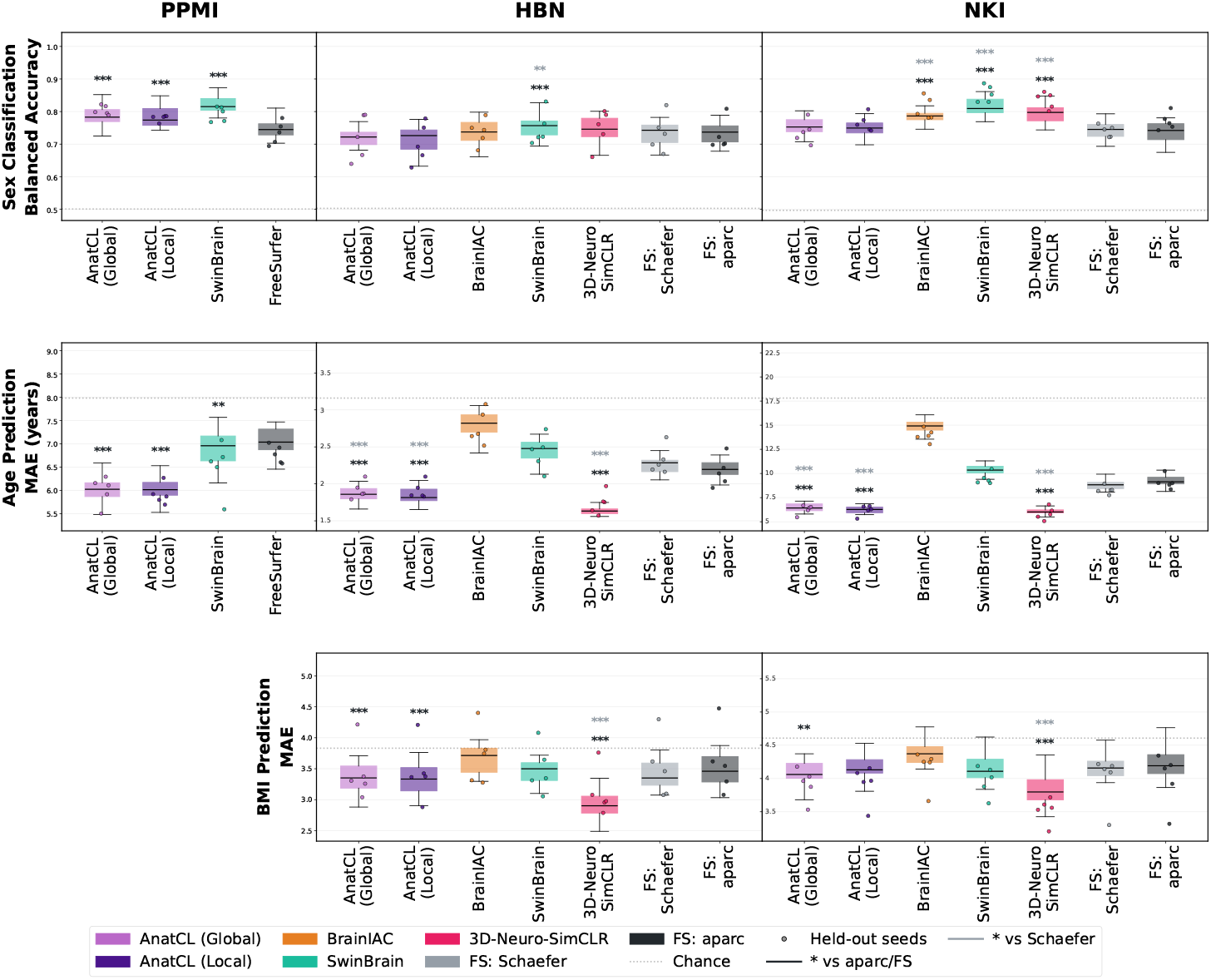
Cross-validation performance at full training size (*α* = 1). Each box displays 5-fold cross-validation performance across 5 random seeds (25 values); overlaid points show held-out test set results for the same 5 seeds. Significance markers correspond to Bonferroni-corrected permutation tests (** *p <* 0.01, ** \**p <* 0.001); dark grey asterisks denote comparisons against FreeSurfer aparc parcellations (FreeSurfer in PPMI), grey asterisks against FS: Schaefer.

#### Sex Classification

In the PPMI dataset, all tested foundation models significantly outperformed the FreeSurfer baseline, with SwinBrain showing the largest improvement. In the HBN dataset, only SwinBrain significantly outperformed the FreeSurfer baselines. In contrast, all models except AnatCL performed better than FreeSurfer baseline on the NKI dataset.

#### Age Regression

In the PPMI dataset, both AnatCL descriptors and SwinBrain outperformed FreeSurfer, with AnatCL achieving the lowest MAE. For both HBN and NKI, AnatCL and 3D-Neuro-SimCLR outperformed the two FreeSurfer baselines, whereas BrainIAC and SwinBrain did not; among all methods, 3D-Neuro-SimCLR produced the lowest MAE on both datasets. Across all datasets, AnatCL consistently outperformed FreeSurfer. While 3D-Neuro-SimCLR was not evaluated on PPMI, it outperformed both baselines on HBN and NKI.

#### BMI Regression

BMI was the most challenging task. 3D-Neuro-SimCLR was the only model to significantly outperform both FreeSurfer baselines on both datasets, achieving the lowest MAE on both HBN (3.11 vs. 3.51 for the strongest FreeSurfer baseline) and NKI (3.52 vs. 3.99). AnatCL significantly outperformed the aparc baseline on both datasets but did not reliably surpass the Schaefer baseline. BrainIAC and SwinBrain did not consistently outperform FreeSurfer. Thus, only 3D-Neuro-SimCLR extracted related signal that reliably exceeded morphometric features.

### 3.2 Foundation model representations differ in alignment with conventional cortical measures

Across all three datasets, we examined correlations between foundation model features and FreeSurfer cortical measures (Fig. 2). AnatCL features—both global and local descriptors—show the strongest overall correlations with FreeSurfer, particularly in HBN and NKI (mean |*r*| ≈ 0.24–0.29). SwinBrain and 3D-Neuro-SimCLR follow at comparable levels, while BrainIAC shows the weakest overall correlations in both cohorts. In PPMI, SwinBrain shows the highest FreeSurfer correlation (|*r*| *≈* 0.21), with AnatCL moderately correlated

**Fig. 2:**
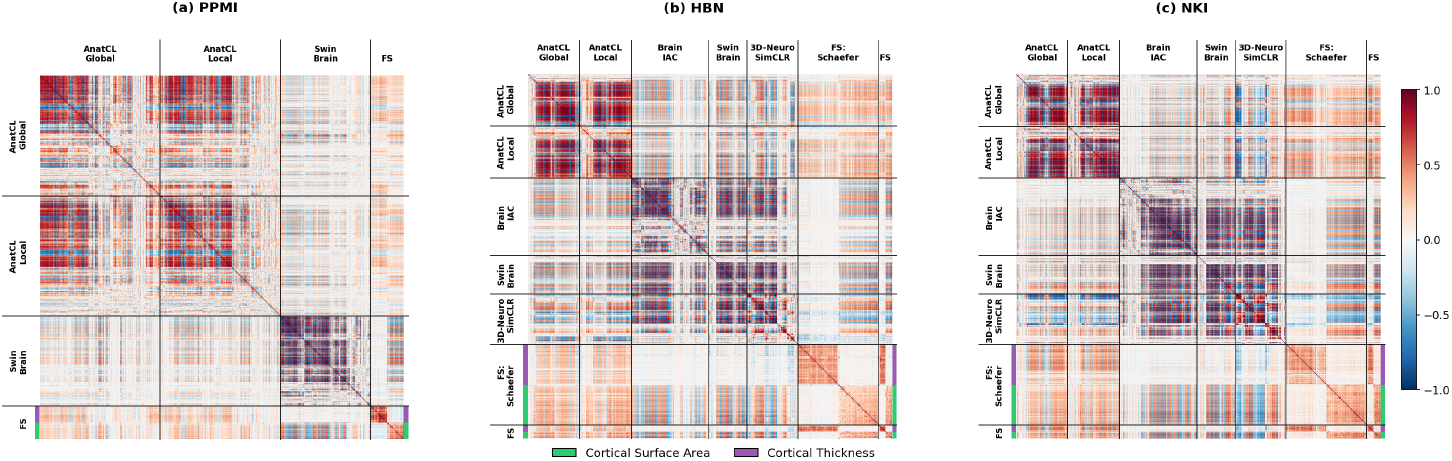
Cross-model correlation matrices showing Pearson correlations between feature dimensions across all model pairs. (a) PPMI, (b) HBN, (c) NKI. Features within each model are reordered by hierarchical clustering (Ward’s method); FreeSurfer features are color-coded by type (green: cortical surface area, purple: cortical thickness).

### 3.3 Results were stable across pre-processing pipelines

Performance of BrainIAC and SwinBrain was consistent across all three preprocessing pipelines (Fig. 3), with no systematic advantage observed for TurboPrep over the alternative pipelines. This suggests that harmonizing preprocessing with TurboPrep did not introduce bias in favor of 3D-Neuro-SimCLR, although neither it nor AnatCL was evaluated under alternative pipelines.

**Fig. 3:**
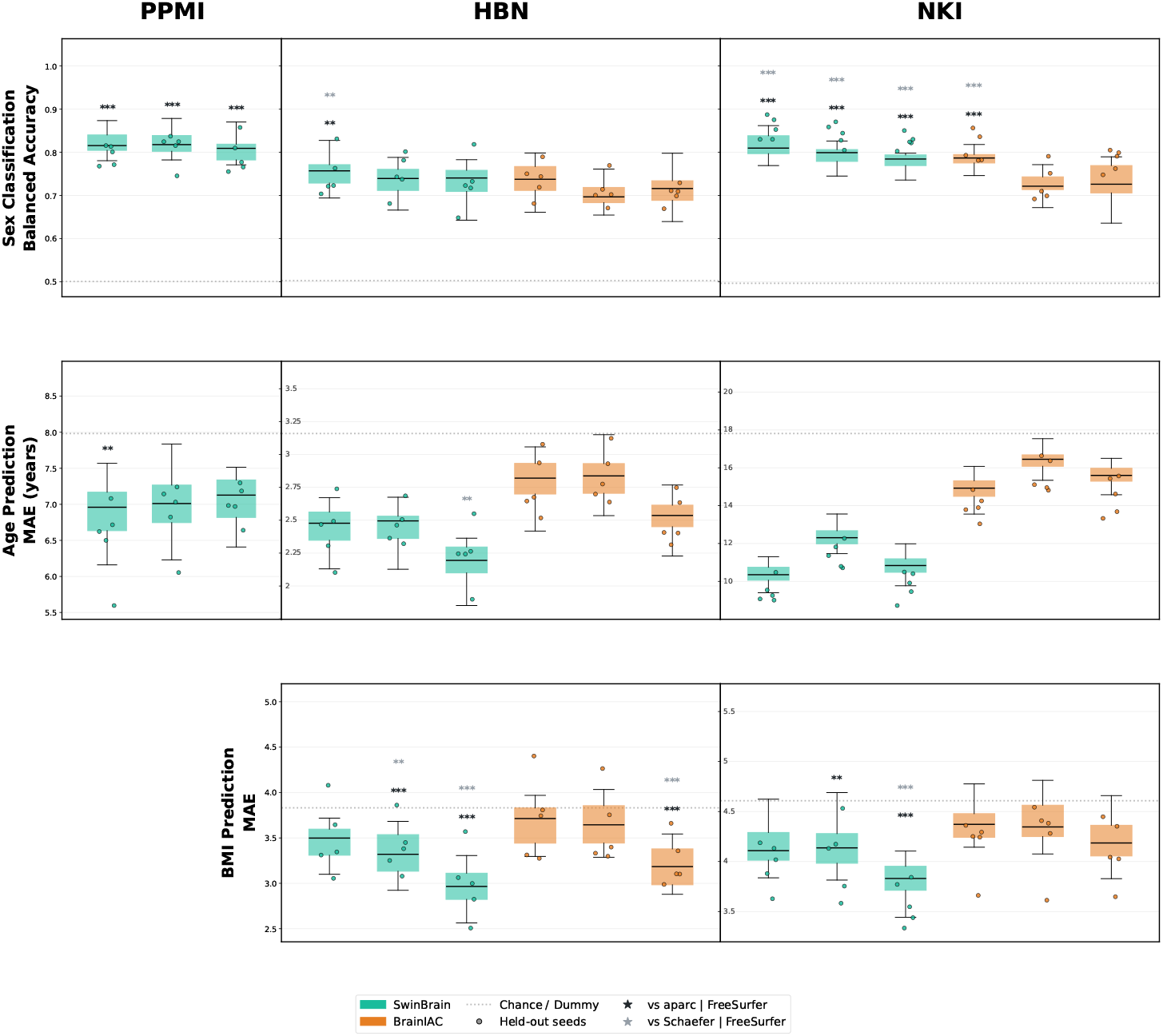
Cross-validation performance of BrainIAC and SwinBrain under three preprocessing pipelines: TurboPrep (left), T1-template registration (middle), and BrainIAC reference template (right). (|*r*| ≈ 0.18). Decomposing these correlations by cortical measure reveals a consistent distinction: SwinBrain, BrainIAC, and 3D-Neuro-SimCLR correlate almost exclusively with cortical surface area (mean |*r*| *≈* 0.33–0.37) rather than thickness (mean |*r*| *≈* 0.06–0.12), whereas AnatCL correlates comparably with both (mean |*r*| *≈* 0.25 for each).

## 4 Limitations

Our evaluation is subject to several limitations. First, the models were employed exclusively as frozen feature extractors; neither fine-tuning nor post-training was conducted. Such adaptation procedures can substantially alter models’ relative performance profiles, and evaluating them falls outside the scope of this benchmark. Second, we compared foundation models against morphometric baselines but not against general-purpose vision encoders, such as a ViT or ResNet pretrained on ImageNet or on web-scale image data, whose transferability to medical imaging tasks has been found to vary across models [9] and which we did not evaluate here. Third, models were trained and evaluated separately within each dataset, so our results characterize performance across three populations rather than generalization across them; cross-dataset and leave-one-site-out evaluation remain future work. Fourth, AnatCL was preprocessed with CAT12 rather than TurboPrep and was not included in our preprocessing comparison. Its features were averaged across five-fold checkpoints, so its advantage may partly reflect pipeline and ensembling rather than representation quality alone. AnatCL was also pretrained with alignment to cortical thickness and surface area, the same measures used in our morphometric baseline. BrainFMBench is designed to accept new models and evaluation settings through pull requests, and we expect these gaps to be addressed as the benchmark grows.

## 5 Conclusion

AnatCL and 3D-Neuro-SimCLR outperformed FreeSurfer-derived features on most tasks and datasets. Notably, both models rely on the lightweight ResNet-18 backbone. This is consistent with, but does not isolate, an effect of architectural inductive bias: the two transformer models also differ from the ResNet-18 models in pretraining data and objective, so architecture cannot be separated from these factors here. In contrast, vision transformer models such as BrainIAC and SwinBrain did not reliably perform better than baseline. These findings were robust to preprocessing choice, as results remained consistent across alternative pipelines applied to BrainIAC and SwinBrain. One reason explaining AnatCL’s performance may be that its pretraining aligns representations with anatomical features including cortical thickness, gray matter volume and surface area, consistent with Fig. 2, in which it is the only model correlating comparably with both thickness and surface area. Similarly, 3D-Neuro-SimCLR may benefit from being pretrained at scale on T1w scans alone, matching the modality of our evaluation datasets. BrainIAC, by contrast, was pretrained on heterogeneous data spanning multiple sequences and clinical conditions, and SwinBrain expects input from three contrasts, which we approximate here by duplicating the T1w volume.

New models can be readily evaluated by submitting a pull request to the BrainFMBench GitHub repository, creating a living resource for the neuroimaging community that enables standardized comparisons of emerging brain structural MRI foundation models as they become available.

## Acknowledgments

Data used in the preparation of this article was obtained on 2025-09-12 from the Parkinson’s Progression Markers Initiative (PPMI) database (www.ppmi-info.org/access-data-specimens/download-data), RRID:SCR_006431. For up-to-date information on the study, visit www.ppmi-info.org. PPMI, a public-private partnership, is funded by the Michael J. Fox Foundation for Parkinson’s Research and funding partners. HBN and NKI data were obtained from the Reproducible Brain Charts (RBC) initiative [16]. This research was enabled in part by support provided by Calcul Québec and the Digital Research Alliance of Canada (alliancecan.ca).

## Disclosure of Interests

The authors have no competing interests to declare that are relevant to the content of this article.

## Footnotes

8 https://github.com/LemuelPuglisi/turboprep

## Notes

### Competing Interest Statement

The authors have declared no competing interest.

### Summary of Updates

Final updated version of the paper.

